# Effects of berberine on growth performance, intestinal microbial, SCFAs, and immunity for Ira rabbits

**DOI:** 10.1101/2023.11.14.567010

**Authors:** Jianing Lu, Xiaoxing Ye, Xinghui Jiang, Mingming Gu, Zhiyi Ma, Qianfu Gan

## Abstract

Berberine (BBR), recognized for its anti-inflammatory and bactericidal properties, has been extensively studied for its effects on mammalian gut microbiota. This study specifically addresses the need for more research on the regulatory effects of BBR on the gut microbiota of Ira rabbits. To fill this gap, we administered varying concentrations of BBR to weaned Ira rabbits to assess its impact on their growth and gut microbiota. In our experiment, 245 healthy weaned rabbits, aged 33 days, were randomly assigned into five groups. The CG group received a standard diet, while groups I, II, III, and IV were given diets supplemented with BBR at doses of 5 mg/kg, 10 mg/kg, 20 mg/kg, and 40 mg/kg, respectively. A 7-day pre-feeding period was implemented for acclimatization, followed by a 30-day experimental phase. The results revealed that BBR significantly improved the Average Daily Feed Intake (ADFI) and Average Daily Gain (ADG) of the rabbits. Notably, group III showed a significantly higher final weight compared to other groups (*P*<0.05). BBR supplementation also increased serum levels of GSH-Px, SOD, and T-AOC, while decreasing MDA levels compared to the control group (*P*<0.05). It also upregulated pro-inflammatory mediators IL-1β, IL-6, and TNF-α, and downregulated anti-inflammatory mediators IL-10 and TGF-β_1_. Furthermore, BBR treatment led to a significant increase in Short-Chain Fatty Acids (SCFAs), specifically acetic and butyric acids (*P*<0.05). Regarding gut microbiota, BBR significantly enhanced the relative abundance of Bacteroidota and Verrucomicrobiota at the phylum level and reduced Firmicutes (*P*<0.05). At the genus level, there was a significant increase in *Akkermansia* and *Alistipes* and a decrease in *Ruminococcus* (*P*<0.05). Overall, BBR appears to promote the growth of Ira rabbits by enriching beneficial bacteria, modulating inflammatory mediators in the TLR4/NF-κB pathway, and reducing inflammation and oxidative stress. Among the tested dosages, 20 mg/kg BBR had the most substantial impact.

## 1 Introduction

The gut microbiota, comprising trillions of microbial communities including bacteria, archaea, and fungi, plays a vital role in the intestinal health of animals (Xing et al., 2022, Renjia et al., 2018). Its disruption can lead to severe consequences such as microbial imbalance, diarrhea, and even death. Research has highlighted the gut microbiota’s significant influence on farm animals’ growth performance, with studies showing its crucial role in metabolism, immunity, and nutrition (Rothschild et al., 2018, Shokryazdan et al., 2017). Notably, microorganisms within the gut microbiota are essential for immune cell development and function, enhancing the body’s ability to combat external threats (Zhuang et al., 2015, Luo et al., 2013). Immunoglobulin, a key indicator of immune efficacy, has been shown to interact with inflammatory factors, highlighting a vital aspect of the body’s immune response (Zhang et al., 2022, Tang et al., 2017).

In the field of animal husbandry, enhancing animal growth performance, reducing mortality rates, and finding safe, effective feed alternatives are ongoing challenges. Studies have shown that gut microbiota significantly impacts farm animals’ growth, and plant extracts have been identified as beneficial feed additives that improve digestion and boost immune function (Karim et al., 2023, Kuldeep et al., 2015).

Berberine (BBR), a compound derived from traditional Chinese medicines such as Coptis chinensis and Cortex Phellodendri, is known for its diverse pharmacological properties (Chen et al., 2008, Habtemariam and Solomon, 2016). It has been recognized for its ability to lower blood sugar and lipids, alleviate diarrhea, and promote intestinal health. Research on BBR has indicated its potential in enhancing mammalian growth performance, with findings suggesting that BBR supplementation can increase animal feed intake and final body weight, offering economic benefits in animal husbandry (Angela and Alberico, 2015, Póciennikowska et al., 2015). Previous studies have explored the potential of BBR in improving mammalian growth performance (Roudini et al., 2019, Sirotkin et al., 2018, Okanlawon et al., 2020). Recent studies also suggest that BBR may regulate intestinal homeostasis by interacting with the gut microbiota. It appears to enhance intestinal permeability and influence the host’s immune system by altering the balance of bacteria that produce short-chain fatty acids (SCFAs) and other metabolites (Sun et al., 2016, Zhang et al., 2020, Yu et al., 2020, Qin et al., 2018, Sarah et al., 2020). However, the specific mechanisms of BBR, especially its impact on animal growth through microorganisms and SCFAs, require further investigation (Sichen et al., 2021, Yue et al., 2019, Wang et al., 2017).

This study aims to explore the mechanism of berberine in regulating intestinal immunity by influencing gut microbiota. The research will include testing metabolic product content, measuring fecal SCFAs, and sequencing 16S rRNA genes. The study will also seek to identify the optimal dosage of BBR and provide deeper insights into how it regulates intestinal immunity. The outcomes are expected to enhance our understanding of the role of gut microbiota and its metabolites in intestinal immune function and growth performance in meat rabbits, a critical area for the rabbit breeding industry.

## 2 Materials and Methods

### 2.1 Animal and experimental design

Ira rabbits, exhibiting similar weights (0.721±0.088kg) and optimal health, were randomly allocated into five groups, totaling 245 rabbits, each aged 33 days, with seven replicates per group (Divided into 3 raised in upper cages and 4 raised in lower cages). Maintaining a consistent nutritional level of the basal diet, berberine (BBR) was added at concentrations of 0 (CG), 5 (I), 10 (II), 20 (III), and 40 mg/kg (IV), respectively, with a preliminary trial duration of seven days and a formal trial period of 30 days. The composition and nutritional level of the basic feed utilized in the experiment are delineated in Table 1, with a berberine concentration of 97% (Shifang Yuheng Plant Raw Materials Co., Ltd., China).

**Table 1:**
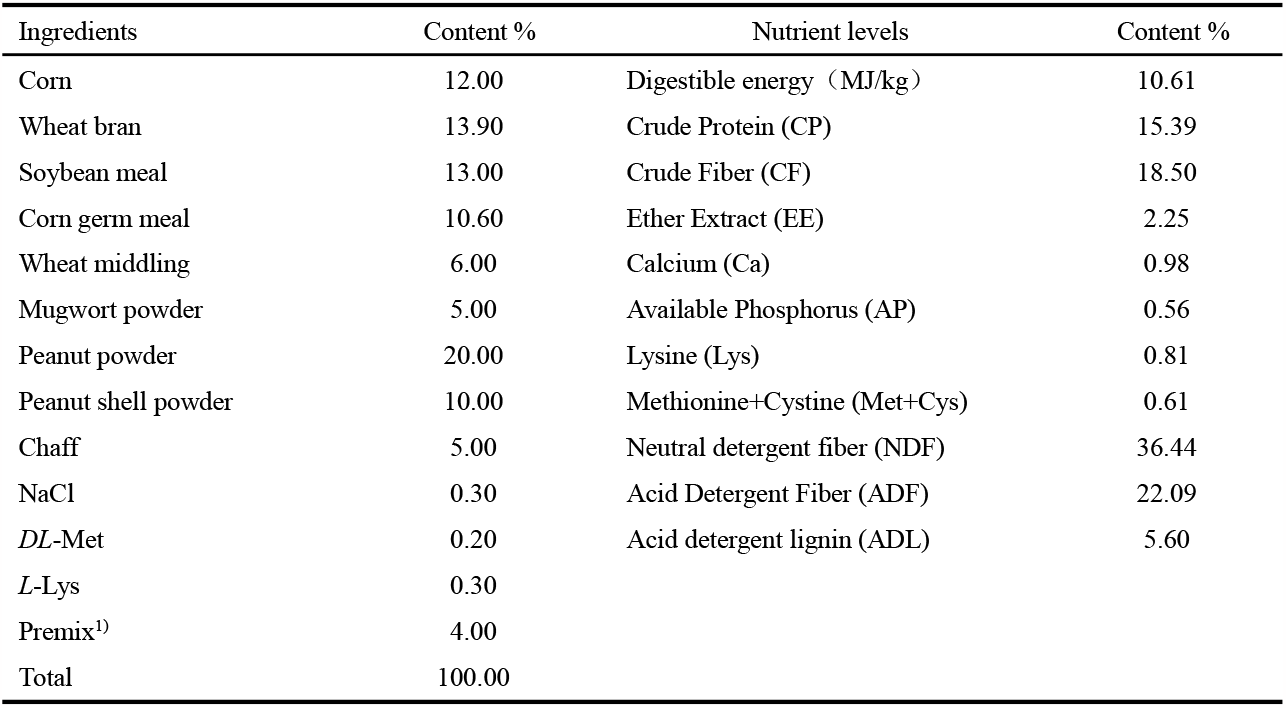
Composition and nutrient levels of basal diet (DM basis)

### 2.2 Sample collection

At the end of the experiment, all Ira rabbits fasted for 12 hours, followed by the random selection of seven meat rabbits from each group, each with a body weight (BW) proximate to the average. After weighing, 5 ml of ear vein blood was collected, centrifuged at 3000 r/min for 10 minutes, and the supernatant was stored at -80°C. Subsequently, the intestine was slaughtered and separated, and sterile EP tubes were used to collect jejunal tissue and its contents, which were stored in liquid nitrogen for the determination of gut microbiota and SCFAs content.

At the beginning and end of the trial period, the surviving rabbits were weighed before morning feeding to calculated the average daily gain (ADG).The formula for ADG is:

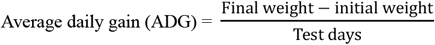

During the experiment, record the number of diarrhea, death, and total feed consumption of Ira rabbits in each group.At the end of the experiment, the ratio of the number of dead rabbits to the number of raised rabbits is the mortality rate, and the ratio of the number of rabbits with diarrhea to the number of raised rabbits is the diarrhea rate.The formula for ADFI is:

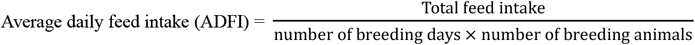

#### 2.2.1 Serum biochemical, immune, and antioxidant indicators

Using Enzyme-Linked Immunosorbent Assay (ELISA) to determine serum immunoglobulin G (IgG), immunoglobulin A (IgA), immunoglobulin M (IgM), interleukin-1β (IL-1β), Interleukin-6 (IL-6), Interleukin-10 (IL-10), Interferon α (TNF-α), Immune Growth Factor β_1_ (TGF-β_1_) Tumor Necrosis Factor γ (IFN-γ) content; Take a blood serum sample to measure blood antioxidant indicators, including total antioxidant capacity (T-AOC), superoxide dismutase (SOD), glutathione peroxidase (GSH-Px), and malondialdehyde (MDA). The specific steps are determined according to the instructions of the kit, which was purchased from Nanjing Jiancheng Biotechnology Research Institute.

#### 2.2.2 Determination of SCFAs content

20 mg of fecal sample were accurately weighed and placed in a 2 mL EP tube. 1 mL of phosphoric acid (0.5% v/v) solution and a small steel ball were added to the EP tube. The samples were ground uniformly, then vortexed for 10 min and ultrasonicated for 5 min. 100 μL of supernatant was moved into 1.5 mL centrifugal tube after the mixture was centrifuged with a speed of 12000 r/min for 10 min at 4°C. 500 μL of MTBE (containing internal standard) solution was added to the centrifugal tube and the mixture was vortexed for 3 min followed by ultrasonicating for 5 min. After that, the mixture was centrifuged with a speed of 12000 r/min for 10 min at 4°C. The supernatant was collected and used for GC-MS/MS analysis (Bianchi et al., 2011, Cesare et al., 2017, KyeongSeog et al., 2022).

#### 2.2.3 16S rRNA gene sequencing analysis

The genomic DNA of jejunal microbial was extracted using cetyltrimethylammonium bromide lysis buffer (CTAB), and the hyper variable region V4-V5 of the bacterial 16S rRNA gene sequencing was amplified using primers 515F(5’-GTGCCAGCMGCCGCGGTAA-3’) and 806R(5’-GGACTACHVGGGTWTCTAAT-3’), through New England Biolabs’ Phusion® High-Fidelity PCR Master Mix with GC Buffer Perform PCR. Using TruSeq® The DNA PCR-Free Sample Preparation Kit kit was used for library construction. The constructed library was quantified by Qubit and Q-PCR, and then sequenced using NovaSeq6000. Use v0.22.0 version of fastp (Chen et al., 2018) to filter the original reads to obtain high-quality reads. The filtering method is to automatically detect and remove joint sequences; Remove reads with an N base amount of 15 or more; Remove reads that account for over 50% of low-quality bases (mass value ≤ 20); Delete items with an average mass of less than 20 within the 4-base window interval; Delete the polyG at the end; Delete reads with a length of less than 150 bp. Then use FLASH version 1.2.11 (Tanja and Salzberg, 2011) to concatenate the sequence and obtain Clean Tags. The Tags sequence is compared with the species annotation database through vsearch (v.2.22.1) to detect chimeric sequences, and the chimeric sequences are ultimately removed to obtain the final effective tags. Cluster all Effective Tags of all samples, and by default, cluster the sequence with 97% consistency to become Operational Taxonomic Units (OTUs). The alpha and beta diversity indices were calculated using QIIME (v.1.9.1). Beta diversity was evaluated through principal component analysis (PCA) graphs based on Euclidean distance. Build a Bar chart using R packages to analyze microbial composition (microorganisms) based on OTU abundance at the phylum and genus levels. LEfSe analysis uses software LEfSe v.1.1.2 (Bolyen et al., 2019), with a default LDA Score filter value of 4.

### 2.3 Statistical analysis methods

The experimental data was analyzed using SPSS 26.0 software for one-way ANOVA. When there were significant differences in ANOVA, Duncan’s method was used for multiple comparisons. Combined with LEfSe analysis, the differences between samples were expressed as enriched OTUs. The experimental results are expressed as mean ± standard deviation, with *P*<0.05 indicating significant differences and *P*<0.01 indicating extremely significant differences.

## 3 Results

### 3.1 Growth performance

The concentration of Berberine (BBR) in the diet significantly enhanced the growth performance of Ira rabbits. Notably, Body Weight (BW), Average Daily Gain (ADG), and Average Daily Feed Intake (ADFI) showed marked improvements (*P*<0.05, Figure 1). Specifically, compared to the Control Group (CG), Groups I, II, III, and IV exhibited BW increases of 2.41%, 6.69%, 12.61%, and 8.42%, respectively (*P*<0.05). Similarly, ADG and ADFI in these groups improved by 10.26%, 21.43%, 23.78%, and 15.70%; and 8.27%, 12.49%, 16.28%, and 9.28%, respectively, compared to CG (*P*<0.05). Moreover, BBR treatment notably reduced the diarrhea and mortality rates in Ira rabbits, with Group III showing particularly low rates of 10.2% for diarrhea and 4.08% for mortality.

**Figure 1:**
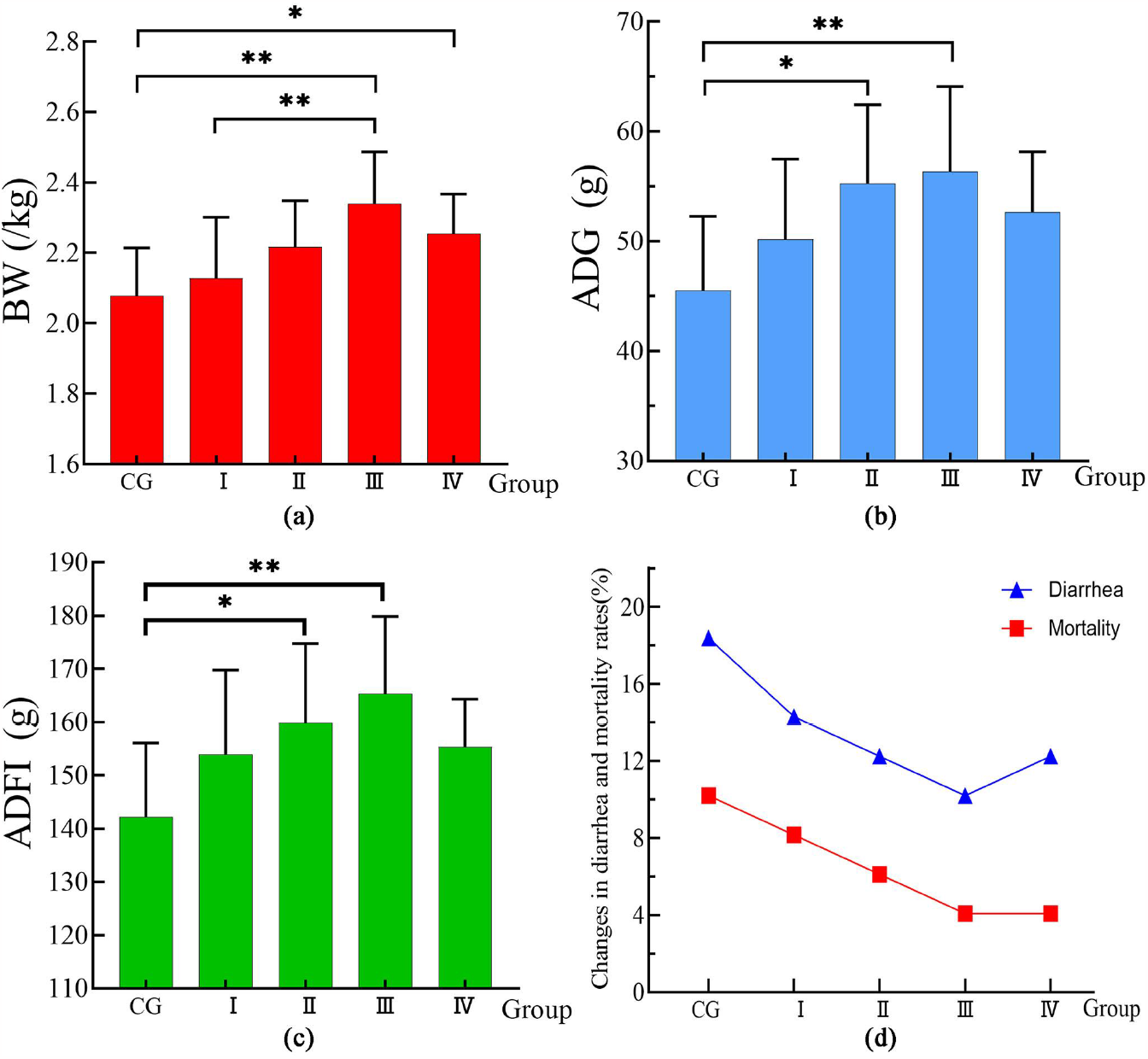
The effect of BBR on the growth performance of Ira rabbits. (a) Body Weight, BW; (b) Average Daily Gain, ADG; (c) Average Daily Feed Intake, ADFI; (d) Diarrhea rate and mortality rate.*P < 0.05, **P < 0.01.

### 3.2 Antioxidant indicators

BBR’s impact on serum antioxidant activity in Ira rabbits is shown in Figure 2. Compared to the CG, BBR-treated rabbits showed a significant decrease in serum Malondialdehyde (MDA) concentration (*P*<0.01), while levels of Glutathione Peroxidase (GSH-Px), Superoxide Dismutase (SOD), and Total Antioxidant Capacity (T-AOC) were significantly higher (*P*<0.01).

**Figure 2:**
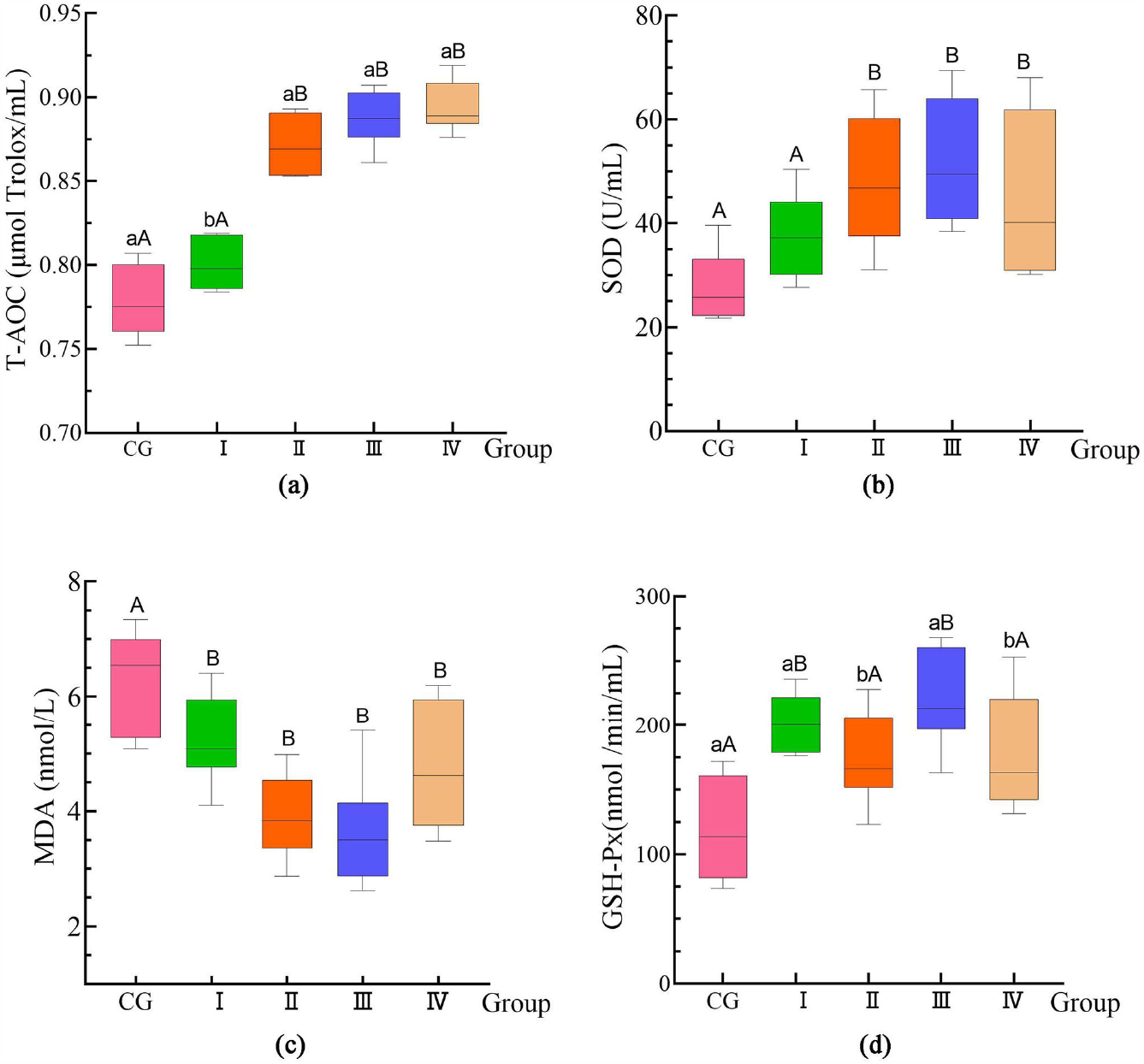
The effect of BBR on the antioxidant capacity of Ira rabbits. (a) T-AOC; (b) SOD; (c) MDA; (d) GSH-Px. Values with no letter or the same letter superscripts mean no significant difference (*P*>0.05), while with different small letter superscripts mean significant difference (*P*<0.05), and with different capital letter superscripts mean significant difference (*P*<0.01). The same as below.The following diagram is the same.

### 3.3 Serum immunity

Figure 3 illustrates the effect of BBR on serum immunoglobulin levels in Ira rabbits. In comparison to the CG, the experimental groups showed significant increases in serum levels of IgA, IgG, IgM, IL-10, and TGF-β1 (*P*<0.05). Additionally, there was a highly significant decrease in IL-1β (*P*<0.01), and notable decreases in IL-6 and TNF-α (*P*<0.05).

**Figure 3:**
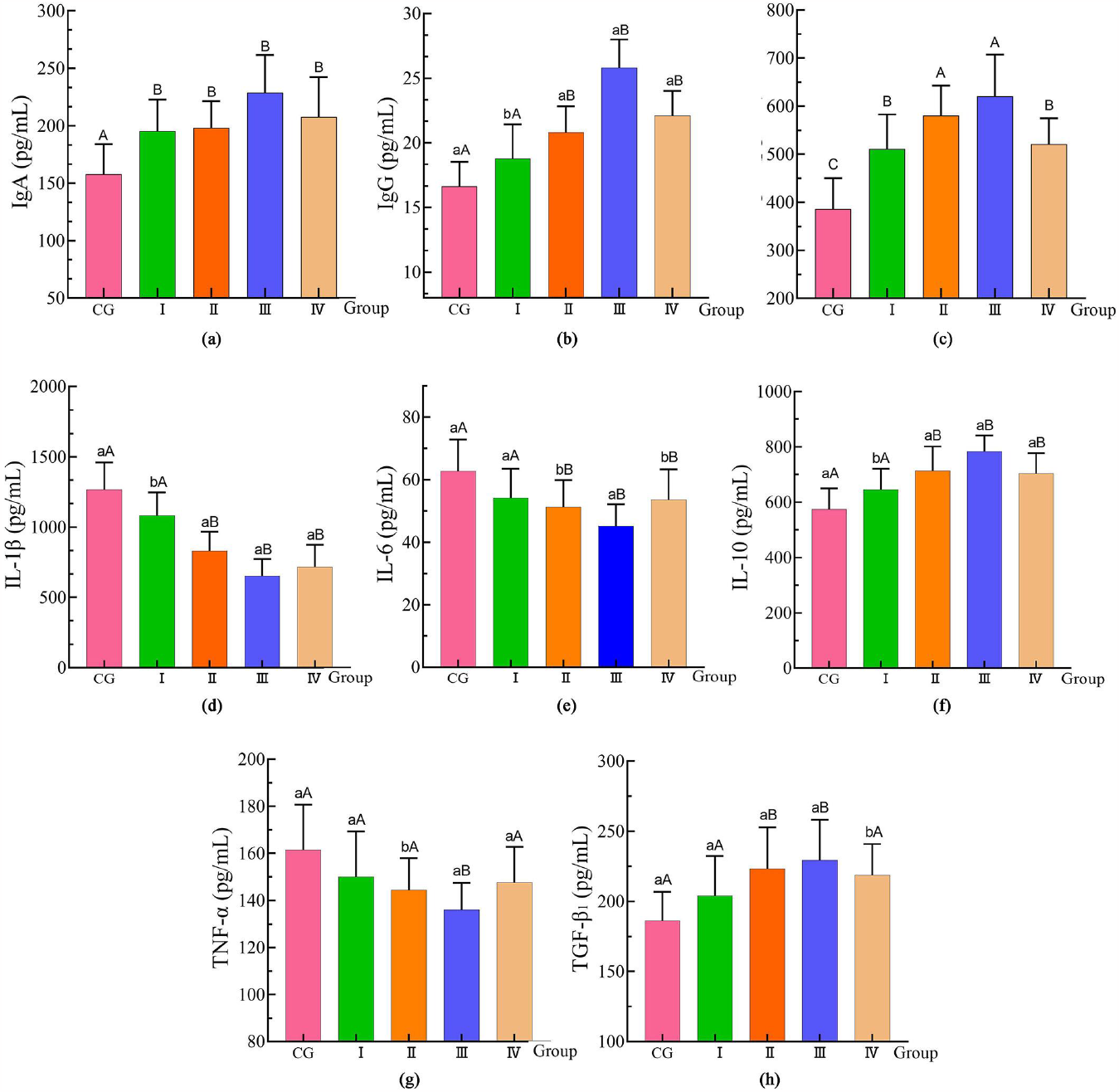
Changes in serum immune function after supplementation with BBR. (a) IgA; (b) IgG; (c) IgM; (d) IL-1β; (e) IL-6; (f) IL-10; (g) TNF-α; (h) TGF-β_1_.

### 3.4 SCFAs

The influence of BBR on SCFA content in Ira rabbits is presented in Figure 4. The levels of acetic acid (AA) and butyric acid (BA) in BBR-treated rabbits were significantly lower compared to the control group (*P*<0.05). Although the level of propionic acid (PA) did not change significantly, a decreasing trend was observed (0.05<*P*<0.1).

**Figure 4:**
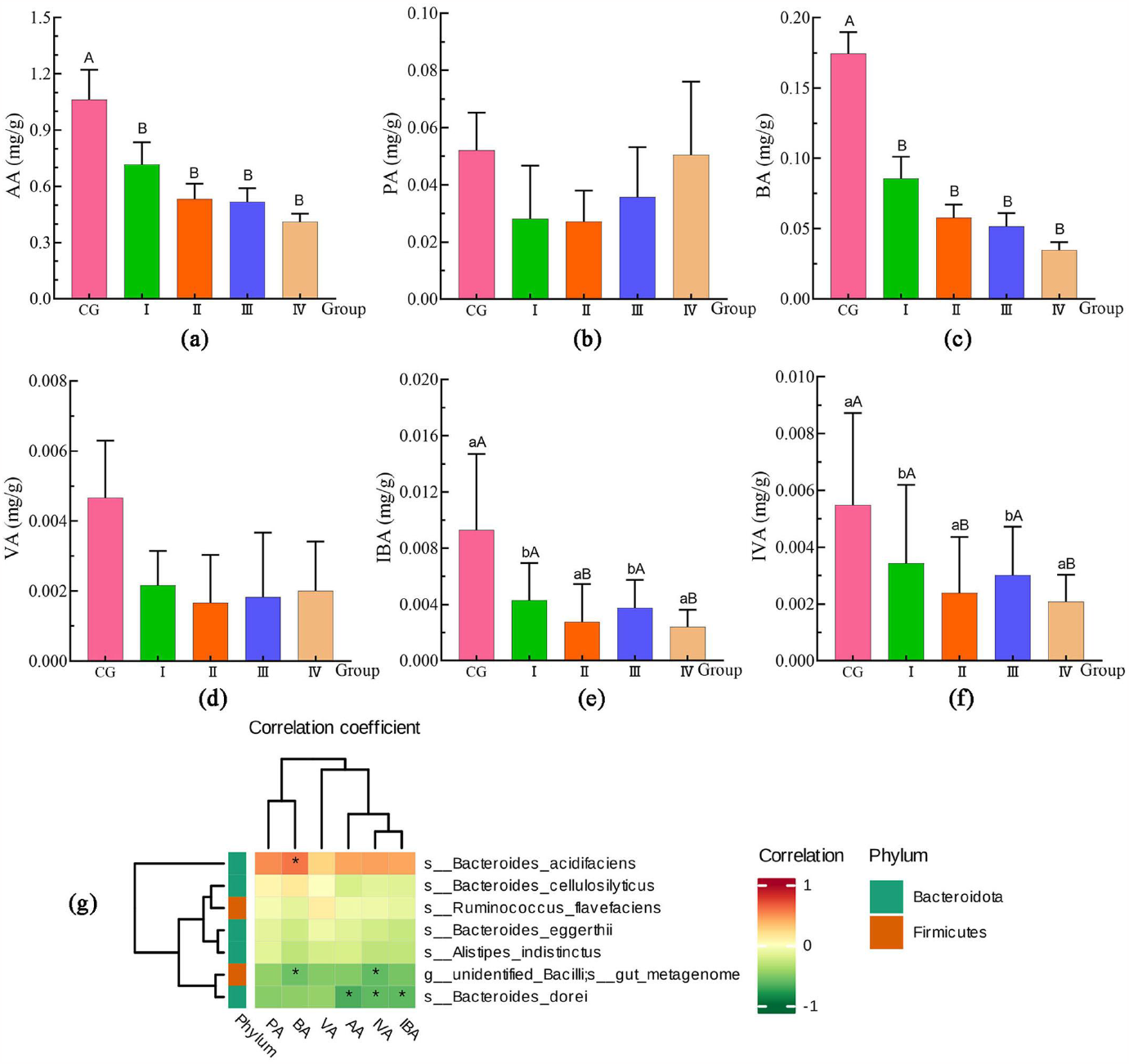
Changes in SCFAs content in the feces of Ira rabbits after supplementing with BBR. (a) AA; (b) PA; (c) BA; (d) VA; (e) IBA; (f) IVA; (g) Correlation analysis between SCFAs and jejunal microbiota.

### 3.5 Intestinal microbiota

The impact of varying BBR supplementation levels on gut microbiota is shown in Figure 5. Between the CG and Groups I, II, III, and IV, there were 6, 10, 16, and 6 differentially enriched Operational Taxonomic Units (OTUs), respectively. BBR supplementation did not significantly alter the diversity index of gut microbiota (Table 2). At the phylum level (Figure 6), Firmicutes and Bacteroidetes were the predominant phyla in each group, with proportions in CG, I, II, III, and IV groups being 77.81%, 66.96%, 65.07%, 65.54%, and 58.25% for Firmicutes, and 17.08%, 24.98%, 24.95%, 27.34%, and 27.20% for Bacteroidetes, respectively. At the genus level, BBR significantly reduced the abundance of *Ruminococcus* and increased the proportions of *Akkermansia* and *Alistipes* (Figure 7).

**Table 2:** The effects of BBR on Alpha diversity index in Ira rabbits.

| Items | CG | I | II | III | IV | P-value |
| --- | --- | --- | --- | --- | --- | --- |
| Chao1 | 1544.13±102.267 | 1658.08±95.293 | 1626.93±97.463 | 1644.73±87.075 | 1629.23±108.92 | 0.324 |
| Simpson | 0.9756±0.0072 | 0.9798±0.01267 | 0.9785±0.0087 | 0.9803±0.0047 | 0.9778±0.0122 | 0.924 |
| shannon | 7.498±0.211 | 7.693±0.496 | 7.657±0.234 | 7.653±0.204 | 7.623±0.255 | 0.668 |
| ACE | 1533.53±122.709 | 1763.87±147.927 | 1671.87±129.139 | 1691.83±126.028 | 1616.28±140.964 | 0.068 |

Data are presented as mean±standard deviation (n=7). In the same row, values with no letter or the same letter superscripts mean no significant difference ( $P>0.05$ ), while with different small letter superscripts mean significant difference ( $P<0.05$ ), and with different capital letter superscripts mean significant difference ( $P<0.01$ ).

**Figure 5:**
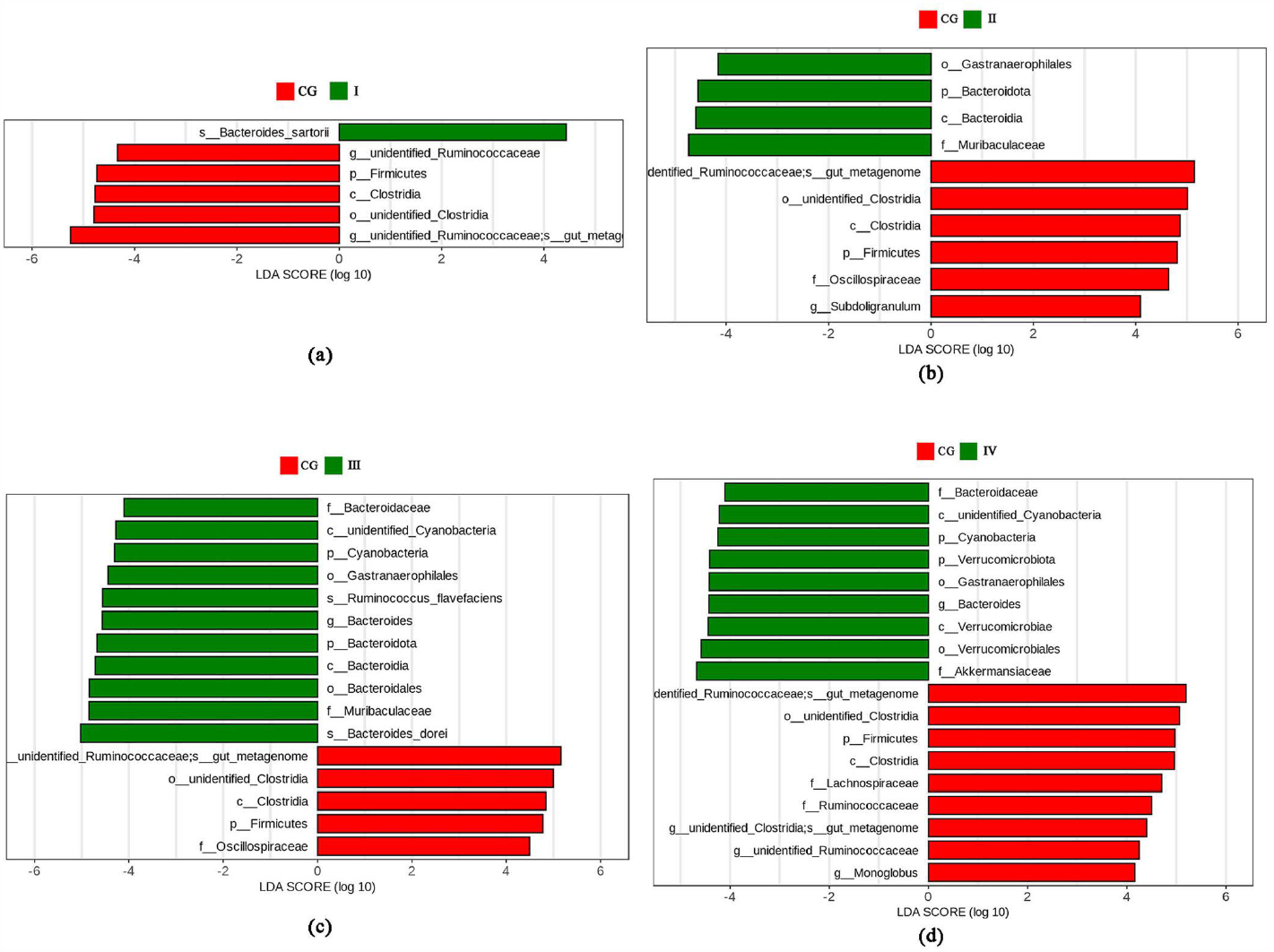
Analysis of variance between groups. (a) CG VS I; (b) CG VS II; (c) CG VS III; (d) CG VS IV.

**Figure 6:**
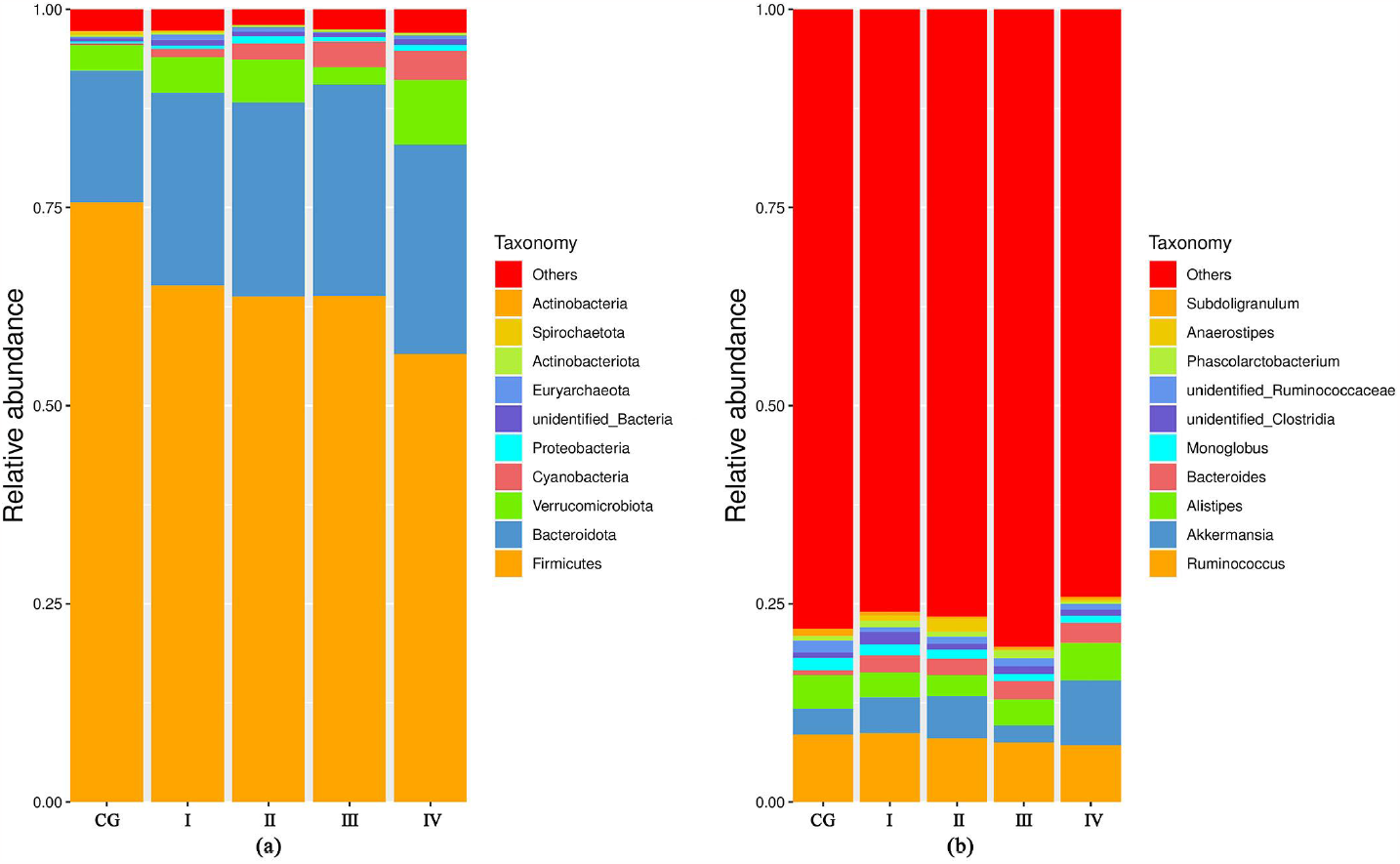
Top 10 phylum level microbiota with relative abundance. (a) phylum level; (b) Genus level.

**Figure 7:**
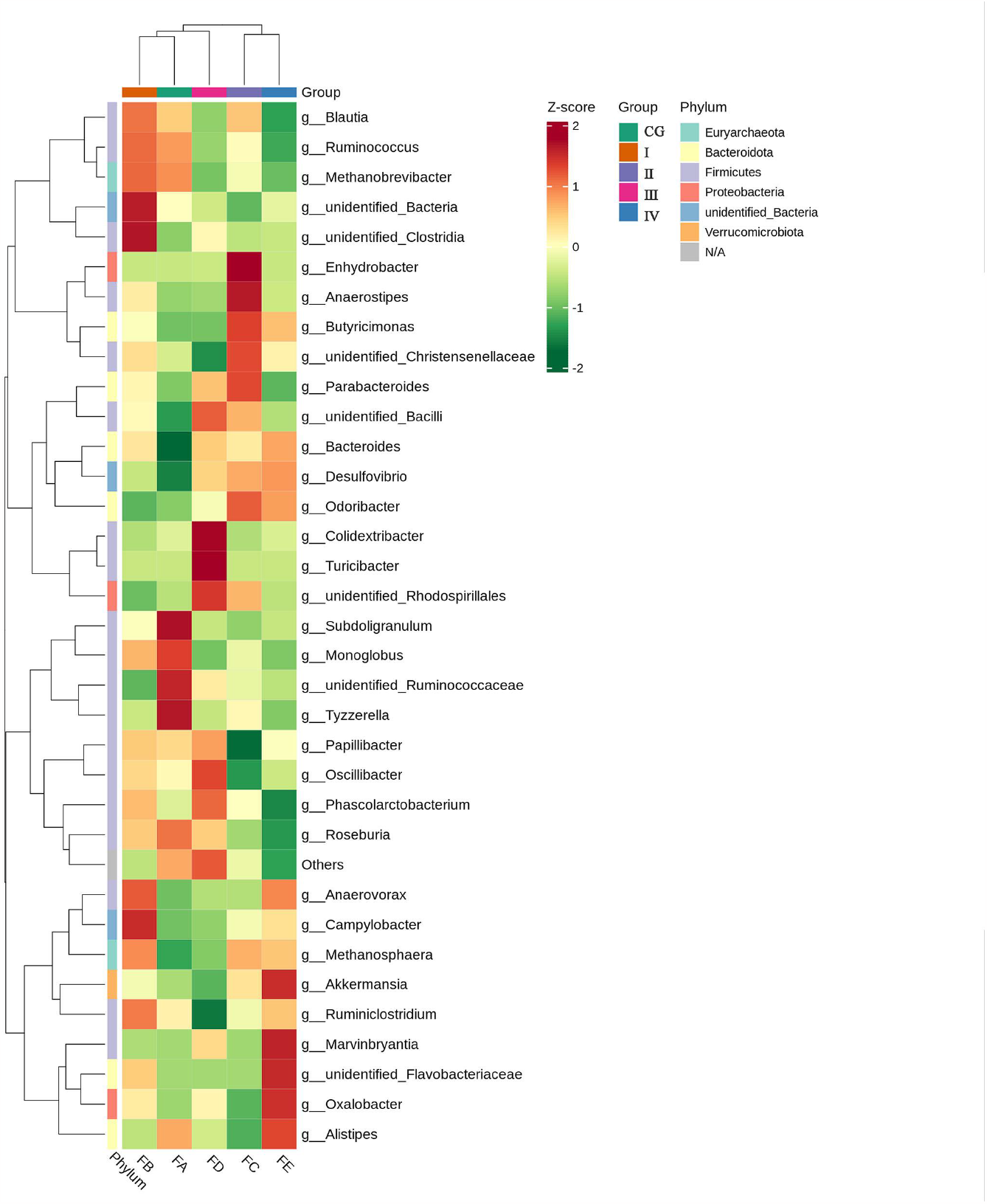
Changes in relative abundance of microorganisms at the genus level.

## 4 Discussion

BBR was chosen as a long-term feed additive for Ira rabbits due to its demonstrated health benefits, including antioxidant, antibacterial, and anti-inflammatory properties (Li et al., 2016, Gong et al., 2017). This study found that BBR supplementation significantly improved the body weight (BW), average daily gain (ADG), and average daily feed intake (ADFI) of Ira rabbits, while reducing diarrhea and mortality rates (Figure 1). The increase in BW may be attributed to BBR’s role in reducing diarrhea and promoting feed intake (Xu et al., 2020, Malekinezhad et al., 2020). The growth-promoting effects of BBR may be attributed to its impact on intestinal health, including enhancing villi height and crypt depth, suppressing inflammatory factors, and boosting intestinal cell absorptive capacity (Scholtz, 2017, Fan *et al*., 2015, Qin et al., 2023, Cui et al., 2020, Zhu et al., 2020, Habtemariam, 2020).

Interestingly, the study revealed that increased serum antioxidants and immunoglobulins correlate with improved animal growth performance (Ma et al., 2021, Maamouri and Salem, 2021). BBR supplementation led to a gradual increase in immunoglobulin content (Figure 3a,b,c), suggesting its role in enhancing animal immune capabilities. However, excessive BBR (40mg/kg) appeared to reduce growth performance, possibly due to its bitter taste affecting palatability and feed intake (Kosalec et al., 2009).

BBR also demonstrated a capacity to enhance antioxidant activities, as evidenced by increased serum Total Antioxidant Capacity (T-AOC), Glutathione Peroxidase (GSH-Px), and Superoxide Dismutase (SOD) activities, and reduced Malondialdehyde (MDA) content (Dongze et al., 2022) (Figure 2). These antioxidant properties are crucial in mitigating oxidative stress and maintaining intestinal health.

Regarding SCFAs, our study observed significant changes in their levels upon BBR supplementation (Figure 4), suggesting BBR’s role in promoting SCFA production by gut bacteria (Wang et al., 2017, Yan et al., 2022, Qin et al., 2023), preventing bacterial and endotoxin translocation into the bloodstream, mitigating inflammatory reactions and immune stress, and averting metabolic disorders (Li et al., 2016; Nilsson et al., 2010; Evans et al., 2010).

The gut microbiota, an integral part of the gastrointestinal system, plays an essential role in the digestion, development, metabolism, and immunity of rabbits (Huang et al., 2021b). The study also highlighted the impact of BBR on gut microbiota composition. BBR supplementation altered the relative abundance of key bacterial phyla and genera, including *Firmicutes, Bacteroidetes*, and *Verrucomicrobiota* (Mattioli et al., 2019, Li et al., 2018, Jin et al., 2018; Figure 5-7). The symbiotic relationship between Bacteroidetes and Firmicutes, key phyla of intestinal bacteria, is crucial for energy absorption or storage in the host. Fluctuations in their populations, such as a decrease in Bacteroidetes or an increase in Firmicutes, are associated with weight gain (Lian et al., 2022, Zhao and Zhang, 2016). These changes in microbiota composition are linked to improved growth performance and reduced diarrhea in Ira rabbits.

In addition, we observed significant differences between the experimental group and the CG group in terms of the relative abundance of genus level. Specifically, as the BBR content increases, the relative abundance of *Ruminococcus* in the experimental group significantly decreases, while there was a marked increase in the relative abundance of *Akkermansia* and *Alistipes* (Figure 7). These alterations suggest BBR’s role in modulating the inflammatory response, as increased *Ruminococcus* is often linked to intestinal inflammatory diseases and inflammation (Victoria Carazo Javier et al., 2023). *Alistipes*, recognized for its anti-inflammatory properties and involvement in lipid metabolism by producing acetic acid from glucose metabolism (McIntosh et al., 2018), and Akkermansia, a representative of Verrucomicrobiota and a potential probiotic for metabolic disorders, are significant. The decrease in inflammation-promoting microorganisms due to BBR may explain the reduced diarrhea incidence in this study. Furthermore, Bacillus, known for its resistance to external stressors and spore-producing ability, and Muribaculaceae, associated with non-diarrhea status, along with Lachnospiraceae, commonly found in obese animals (Na et al., 2018), showed an increase following BBR supplementation (Figure 7). These results indicate BBR’s potential in reducing diarrhea and promoting weight gain in Ira rabbits through microbial modulation.

The TLR4/NF-κB signaling pathway, essential in cell proliferation, apoptosis, and inflammatory respons (Yang et al., 2018, Iribarren et al., 2015, Xiong et al., 2017, He et al., 2017, Wu et al., 2017, Keren et al., 2018, Sinagra et al., 2017, Choi et al., 2019, Su et al., 2014, Tang et al., 2021, Min et al., 2020, Chu et al., 2014), is activated by TLR4, a transmembrane protein on intestinal and immune cells, upon recognizing pathogens (Duan et al., 2022). NF-κB activation leads to inflammatory reactions (He et al., 2017) and the release of pro-inflammatory factors (Wu et al., 2017; Keren et al., 2018; Sinagra et al., 2017). Our study observed a reduction in serum IL-1β, IL-6, and TNF-α levels following BBR supplementation (Figure 3,d,e,g), indicating its potential in inhibiting pro-inflammatory factor production. The suppression of inflammatory factor expression, known to reduce inflammation, is achieved by inhibiting the NF-κB pathway (Choi et al., 2019; Su et al., 2014). BBR’s regulation of NF-κB and MAPK pathways in mitigating intestinal injury, immune suppression, and oxidative stress, thereby maintaining intestinal health in piglets (Tang et al., 2021). We propose that BBR reduces inflammatory response and oxidative stress by inhibiting the TLR4/NF-κB pathway.

In conclusion, BBR positively influences the growth performance and immune function of Ira rabbits by modulating gut microbiota, SCFAs, and inflammatory pathways. The optimal BBR dosage of 20mg/kg (Group III) yielded the best overall results. These findings underscore BBR’s potential as a feed additive, enhancing intestinal immune function and alleviating diarrhea.

## 5 Conclusions

The findings of our study indicate that supplementing Ira rabbits with BBR significantly enhances their growth performance, with the most favorable effects observed at a dosage of 20mg/kg. Additionally, BBR was found to alter the composition of the gut microbiota and bolster the immune capacity of the rabbits. Our research offers insights into the potential mechanisms by which modulating the gut microbiota can reduce diarrhea rates and further improve rabbit growth performance. These results not only deepen our understanding of BBR’s role but also open avenues for future research aimed at optimizing the growth performance of Ira rabbits.

## 6 Acknowledgments

We sincerely thank Mr. Chen Zhoulin, the manager of the commercial Ira rabbit farm, for providing the opportunity to collect samples and conduct phenotypic measurements. Thank you very much to all laboratory colleagues who helped measure data and test indicators during the experiment.

## 7 Authors’ Contributions

JN contributed to conception and design of the study, performed the experiments, revised the manuscript; XX analyzed the data, wrote and revised the manuscript; XH performed the experiments, analyzed the data, and revised the manuscript; MM and ZY performed the experiments; QF conceived and designed the experiments, supervised the experiment progress, revised the manuscript. All authors read and approved the final manuscript.

## 9 Tables and Figures

